# Structural basis for ligand-induced inactivation of protein tyrosine receptor type Z (PTPRZ): Physiological relevance of head-to-toe RPTP dimerization

**DOI:** 10.1101/636423

**Authors:** Akihiro Fujikawa, Hajime Sugawara, Naomi Tanga, Kentaro Ishii, Kazuya Kuboyama, Susumu Uchiyama, Ryoko Suzuki, Masaharu Noda

**Affiliations:** Division of Molecular Neurobiology, National Institute for Basic Biology (NIBB), 5-1 Higashiyama, Myodaiji-cho, Okazaki, Aichi 444-8787, Japan; Asubio Pharma Co., Ltd., 6-4-3 Minatojima-Minamimachi, Chuo-ku, Kobe, Hyogo, 650-0047, Japan; School of Life Science, The Graduate University for Advanced Studies (SOKENDAI), 5-1 Higashiyama, Myodaiji-cho, Okazaki, Aichi 444-8787, Japan; Exploratory Research Center on Life and Living Systems (ExCELLS), National Institutes of Natural Sciences, 5-1 Higashiyama, Myodaiji-cho, Okazaki 444-8787, Japan; Department of Biotechnology, Graduate School of Engineering, Osaka University, 2-1 Yamadaoka, Suita, Osaka, 565-0871, Japan; Research Center for Cell Biology, Institute of Innovative Research, Tokyo Institute of Technology, 4529 Nagatsuta-cho, Midori-ku, Yokohama, Kanagawa 226-8503, Japan

**Keywords:** cellular signaling, chondroitin sulfate, dimerization, knock-in mouse, mass spectrometry, oligodendrocyte, phosphotyrosine, pleiotrophin, receptor-type tyrosine phosphatase, X-ray crystallography

## Abstract

Protein tyrosine phosphatase receptor type Z (PTPRZ) has two receptor isoforms (PTPRZ-A and -B) containing tandem PTP-D1 and -D2 domains intracellularly, with only D1 being active. Pleiotrophin (PTN) binding to the extracellular region of PTPRZ leads to the inactivation of PTPase, thereby inducing oligodendrocyte precursor cell (OPC) differentiation and myelination in the CNS. However, the mechanisms responsible for the ligand-induced inactivation of PTPRZ remain unclear. We herein revealed that the crystal structure of the intracellular region of PTPRZ (PTPRZ-ICR) showed the “head-to-toe”-type dimer conformation, with D2 masking the catalytic site of D1. Mass spectrometry (MS) revealed that PTPRZ-ICR proteins remained in monomer-dimer equilibrium in aqueous solution, and a substrate-derived inhibitory peptide or competitive inhibitor (SCB4380) specifically bound to the monomer form in a 1:1 stoichiometric ratio, supporting the “head-to-toe dimerization” model for inactivation. A D2 deletion (ΔD2) or dimer interface mutation (DDKK) disrupted dimer formation, while the binding of SCB4380 was maintained. Similar to wild-type PTPRZ-B, monomer-biased PTPRZ-B-ΔD2 and PTPRZ-B-DDKK mutants efficiently dephosphorylated p190RhoGAP at Tyr-1105 when co-expressed in BHK-21 cells. The catalytic activities of these mutants were not suppressed by a treatment with PTN, but were inhibited by the cell-permeable PTPase inhibitor NAZ2329. The PTN treatment did not enhance OPC differentiation in primary cultured glial cells prepared from ΔD2 or catalytically-inactive CS mutant knock-in mice. Our results indicate that PTN-induced PTPRZ inactivation is attained by dimer formation of the intracellular tandem PTP domains in the head-to-toe configuration, which is physiologically relevant to the control of OPC differentiation *in vivo*.

---

The tyrosine phosphorylation of cellular proteins in higher eukaryotes serves as an important signal transduction system in various cellular events, including cell proliferation, migration, and differentiation (1). This phosphorylation is reversibly controlled by protein tyrosine kinases (PTKs) and protein tyrosine phosphatases (PTPs). The regulatory mechanisms of receptor PTKs (RPTKs) have been extensively examined; the ligand-induced dimerization of RPTKs leads to the transphosphorylation of key tyrosine residues in the activation loop, resulting in the full activation of its catalytic activity (2).

In contrast to RPTKs, the dimerization-induced inactivation model of receptor-type PTP (RPTP) activity has remained controversial. RPTPs have a similar structure comprising a unique extracellular domain, single transmembrane domain, and one or two intracellular PTP domains. In the latter case with two PTP domains, the membrane proximal PTP domain (D1) exhibits PTP activity, whereas the distal PTP domain (D2) has little or no catalytic activity (3). In the late 1990s, the dimerization-induced inactivation mechanism of RPTP was initially suggested by structural studies on the D1 of PTPRA/PTPα (4,5). The crystal structure of PTPRA-D1 indicated a dimeric assembly in which the N-terminal helix-turn-helix wedge motif occluded the active site in the catalytic domain of the interacting counterpart. However, subsequent structural studies on the cytoplasmic tandem D1D2 segments of PTPRF/LAR (6), PTPRC/CD45 (7), and PTPRG/PTPγ (8) showed that the orientations of the D1 and D2 domains were highly conserved and incompatible with the “inhibitory wedge model” due to a steric clash. In 2009, Barr AJ et al. reported a novel “head-to-toe dimer” structure for the cytoplasmic D1D2 of PTPRG, in which D1 of one molecule interacted with D2 of the other molecule (counterpart) and vice versa (8). However, the authors stated that “*future studies are necessary to establish this molecular mechanism* in vivo *as a form of inhibitory regulation*”.

PTPRZ is a member of the R5 RPTP subfamily together with PTPRG (3). Both molecules structurally resemble each other and contain a carbonic anhydrase (CAH)-like domain and fibronectin type III-like domain extracellularly as well as two tyrosine phosphatase domains intracellularly. Three splicing variants of PTPRZ are known: PTPRZ-A, a full-length receptor form; PTPRZ-B, a short receptor form with a deletion in the extracellular region; and PTPRZ-S (also called phosphacan), a secretory form (9). These receptor isoforms have been further subdivided into two sub types; conventional PTPRZ-A and -B and exon 16-deleted PTPRZ-A_Δex16_ and -B_Δex16_, respectively (10).

All three isoforms expressed in the CNS are heavily modified with chondroitin sulfate chains on their extracellular segment (9). The chondroitin sulfate moiety is essential for the diffuse distribution of PTPRZ receptors on the cell membrane, thereby maintaining them in their catalytically active monomer state. Chondroitin sulfate chains are required for achieving the high-affinity binding sites of endogenous inhibitory ligands, such as PTN/heparin-binding growth-associated molecule (HB-GAM), midkine, bFGF, and interleukin-34 (11). Positively charged ligands induce PTPRZ clustering, potentially by neutralizing electrostatic repulsion between negatively charged chondroitin sulfate chains, thereby inactivating the intracellular catalytic activities of PTPRZ receptors through dimerization in living cells (11). In contrast, PTPRG is not modified by chondroitin sulfate chains on the extracellular region, and no extracellular ligands have been identified to date.

PTPRZ is the most abundant RPTP molecule in oligodendrocyte precursor cells (OPCs) (12). It functions as a counterpart of FYN tyrosine kinase and dephosphorylates p190RhoGAP (13). The expression of PTPRZ-A, PTN, and myelin basic protein (MBP) peaks on postnatal day 5 (P5), P10, and P30, respectively, during development. On P10, the amounts of MBP and myelinated axons in the brain were shown to be significantly higher in *Ptprz*-null knockout mice than in wild-type animals (11). Conversely, *Ptn*-knockout mice exhibited delayed myelination (11). Consistent with these *in vivo* findings, oligodendrocytes appear earlier in primary cultures from *Ptprz*-deficient mice than from wild-type mice (12). In addition, PTN promotes thyroid hormone-induced OPC differentiation in glial cells obtained from wild-type mice, but not those from *Ptprz*-null knockout mice (12). These findings indicate that PTN-PTPRZ signaling fine-tunes the timing of oligodendrocyte differentiation and myelination in the developing mouse brain. However, there is currently no experimental evidence to show that the ligand-induced inactivation of RPTP molecules is performed by intracellular dimer formation or that it is relevant to physiological events.

In the present study, we show that the crystal structure of rat PTPRZ has the “head-to-toe dimer” configuration. The results obtained from several distinct methods, including native mass spectrometry (MS) and cell-based assays, support the validity of this inactivation mechanism in the ligand-induced inactivation of PTPRZ in living cells. Furthermore, primary cultured glial cells show its physiological relevance to the mechanisms responsible for OPC differentiation. In the Discussion section, we added overall considerations regarding the dimer interface region in RPTP family members.

## RESULTS

### Crystal structures of tandem PTP domains of rat PTPRZ

We achieved the crystallization of the whole intracellular region of rat PTPRZ (rat PTPRZ-ICR) using the sitting-drop vapor-diffusion method, and succeeded in solving the structure at 3.32 Å resolution (Fig. 1 and Table S1). The structure showed “head-to-toe dimer” formation, as reported for the structure of PTPRG-D1D2 (PDB accession number, 2NLK) (8). Contact sites across the dimer interface are shown in Figure 2A. This interaction involved extensive electrostatic interactions between an electropositive pocket at the active center of D1 in one molecule and an electronegative surface between strands β10–β11 of D2 in the other molecule (see the right panels in Fig. 2Aa and Ab). The approximate area of the solvent-accessible interface between the two molecules was calculated to be 1230 Å^2^ (5.1% of the total surface), and multiple hydrogen bonds and salt bridges were estimated using PDBePISA (Table S2), with two residues of Asp-2178 and Asp-2179 within the loop connecting strands β10–β11 in D2 exhibiting the most extensive interactions (Fig. 2B). These structural features are similar to those reported for the dimer structure of PTPRG (8).

**Figure 1.**
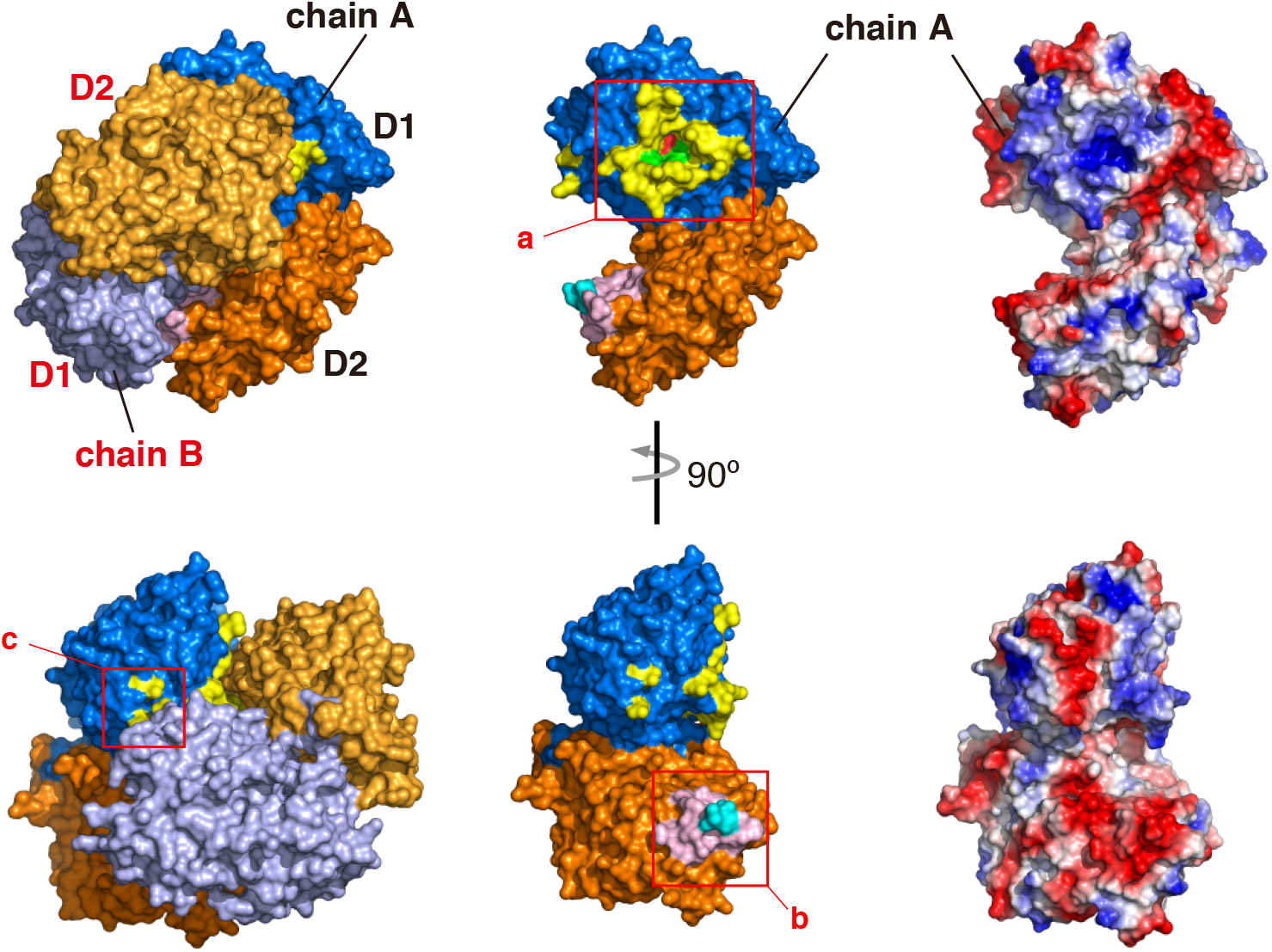
PTPRZ-ICR dimer structure. Surface representations of rat PTPRZ-ICR (PDB, 6J6U) in which the two molecules (chains A and B) formed a “head-to-toe dimer”, with the D1 domain (blue and light purple) of one molecule interacting with the D2 domain of a second molecule (orange and yellow-orange) (left). Chain A extracted from the dimer is shown by surface representations (center) or electrostatic potentials colored red for a negative charge and blue for a positive charge (right). The three interface regions are indicated by square boxes (a to c), and their details are shown in Figure 2A.

**Figure 2.**
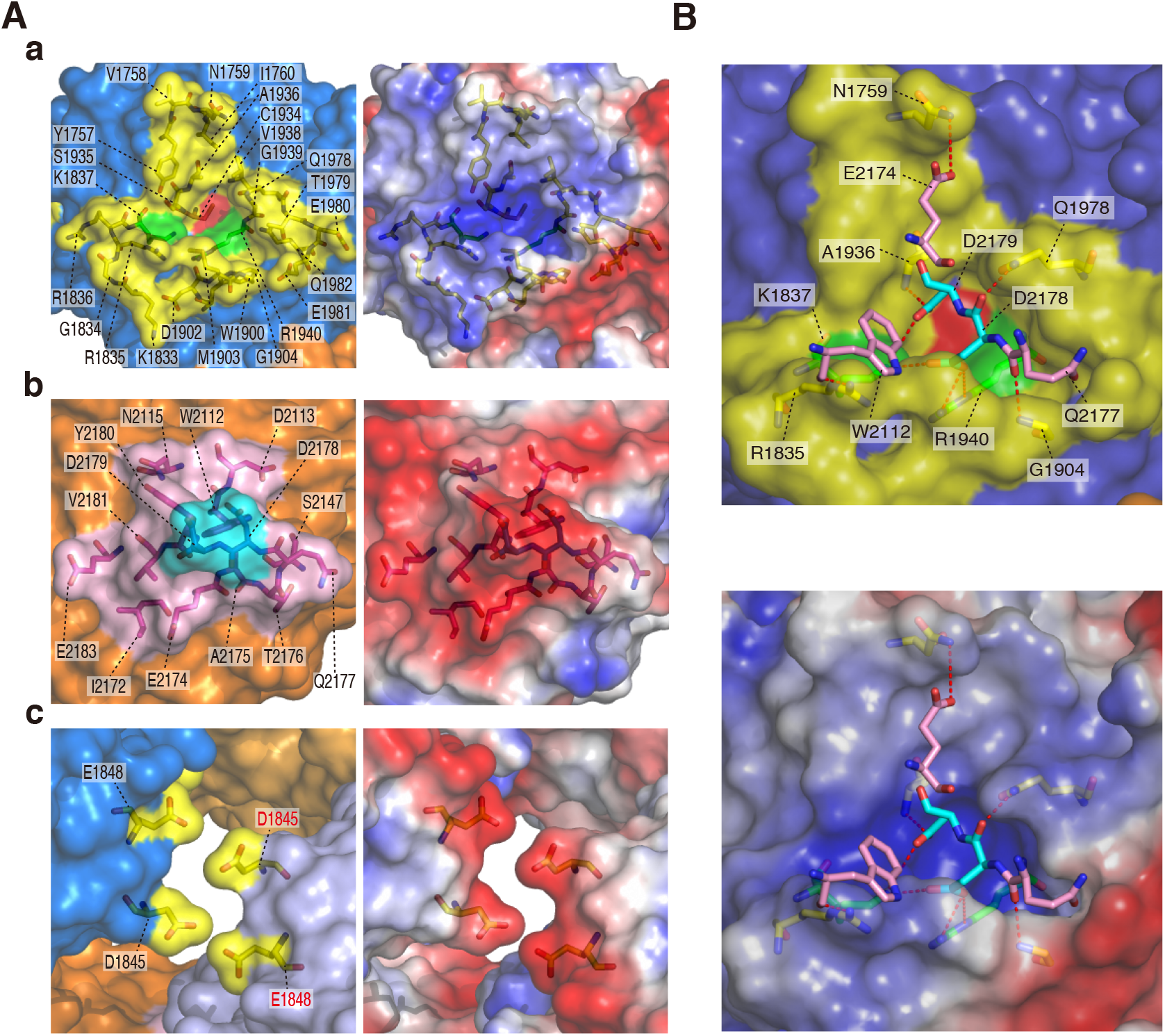
Dimer interface. ***Aa-c***, Amino acid residues forming the dimer interface are shown by a stick model inside the transparent surface (left) and electrostatic potentials (right). Interface regions are colored as in Figure 1. ***B***, Interactions between the D1 and D2 dimer interfaces are shown by red dashed lines (Details are given in supplementary Table S1).

### Masking the active site in D1 with D2 in response to ligand binding

Mass spectrometry (MS) under nondenaturing conditions, referred to as native MS, was used to investigate non-covalent molecular interactions in aqueous solution (14). Native MS revealed that PTPRZ-ICR proteins were in monomer-dimer equilibrium in solution, and dimers were observed in the concentrations higher than 3.2 μM (Fig. 3A). If the “head-to-toe dimer” configuration was present in solution as well as in the crystal structure, substrate binding may only take place in the monomer state. To test this possibility, we used a non-hydrolyzable pTyr mimetic (F2Pmp, phosphonodifluoromethyl phenylalanine)-containing peptide (15) based on the typical substrate motif for PTPRZ (16): The substrate mimetic, F_2_Pmp-GIT1_550–556_, effectively inhibited the catalytic activity of PTPRZ-ICR (Ki = 3.1 μM).^4^ In native MS analyses, F_2_Pmp-GIT1_550–556_ specifically bound to the monomer fraction of PTPRZ-ICR in a 1:1 stoichiometry, while no binding was observed in the dimer fraction (Fig. 3B). Similar results were obtained when mixed with the competitive PTPRZ inhibitor, SCB4380 (Fig. 3C). Notably, increases in F_2_Pmp-GIT1_550–556_- or SCB4380-bound monomers inversely correlated with decreases in the dimer fraction. These results indicated that PTPRZ-ICR continuously shifts between monomer and dimer states, and that the substrate mimic prevents dimer formation when bound to the catalytic pocket in D1.

**Figure 3.**
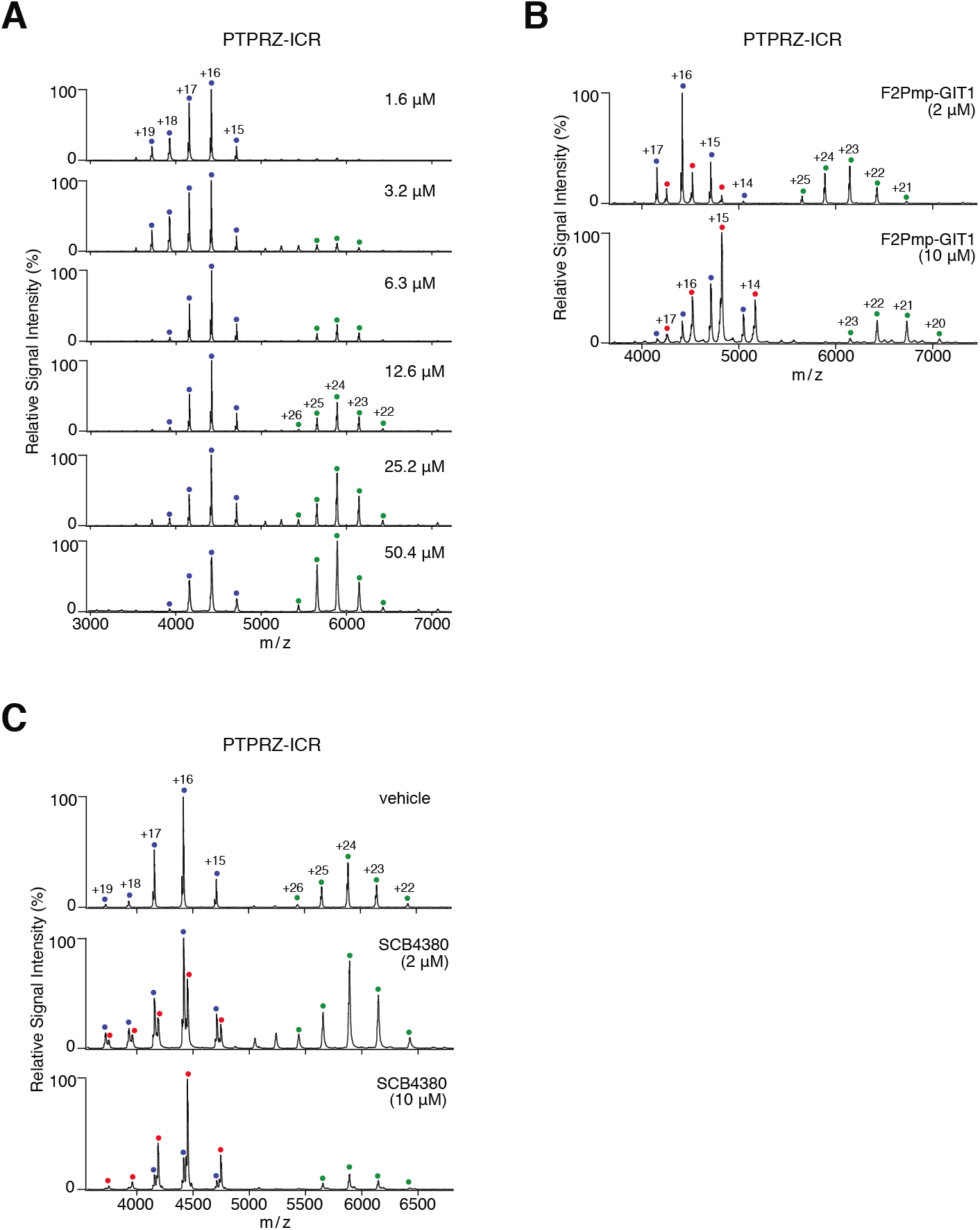
Native MS analyses for the monomer-dimer equilibrium of rat PTPRZ-ICR and inhibitor binding in aqueous solution. ***A***, The monomer-dimer equilibrium of rat PTPRZ-ICR proteins at the indicated concentrations was assessed by MS under non-denaturing conditions. Product ion spectra derived from monomers (corresponding to a molecular mass of 70,639 ± 1 Da) or dimers (141,287 ± 8 Da) are indicated by blue and green dots with charge states, respectively. ***B,C***, PTPRZ-ICR proteins (12.6 μM) were mixed with a substrate-mimetic inhibitor peptide, F_2_Pmp-GIT1 (B) or small competitive inhibitor molecule, SCB4380 (*C*) at the indicated concentrations, and then subjected to the MS analysis. Peaks corresponding to PTPRZ-ICR monomers complexed with SCB4380 (71,178 ± 2 Da) or F_2_Pmp-GIT1 (72,322 ± 1 Da) are indicated by red dots with charge states. MS peaks corresponding to PTPRZ-ICR dimers complexed with SCB4380 or F_2_Pmp-GIT1 were not observed. Masses were estimated from their parent ion peaks, and shown as the mean ± standard deviation (SD) of three measurements.

We then produced two PTPRZ-ICR mutant proteins: one (DDKK mutant) in which Asp-2178 and Asp-2179 were substituted with Lys residues to test its dimerization ability in solution; the other was the D2 domain-deleted form (ΔD2 mutant) to examine its blocking ability of the catalytic site of D1. ΔD2 and DDKK mutant enzymes exhibited similar PTPase activities, and their IC_50_ values for SCB4380 were similar to that of wild-type PTPRZ-ICR (Fig. S1). Both mutant proteins remained mostly as monomers within the concentrations tested and bound SCB4380 in a 1:1 stoichiometry in native MS (Fig. 4), indicating that SCB4380 did not bind to D2. These results strengthened our view that “head-to-toe dimer” formation underlies the ligand-induced inactivation of PTPRZ. These results were summarized in Figure 5.

**Figure 4.**
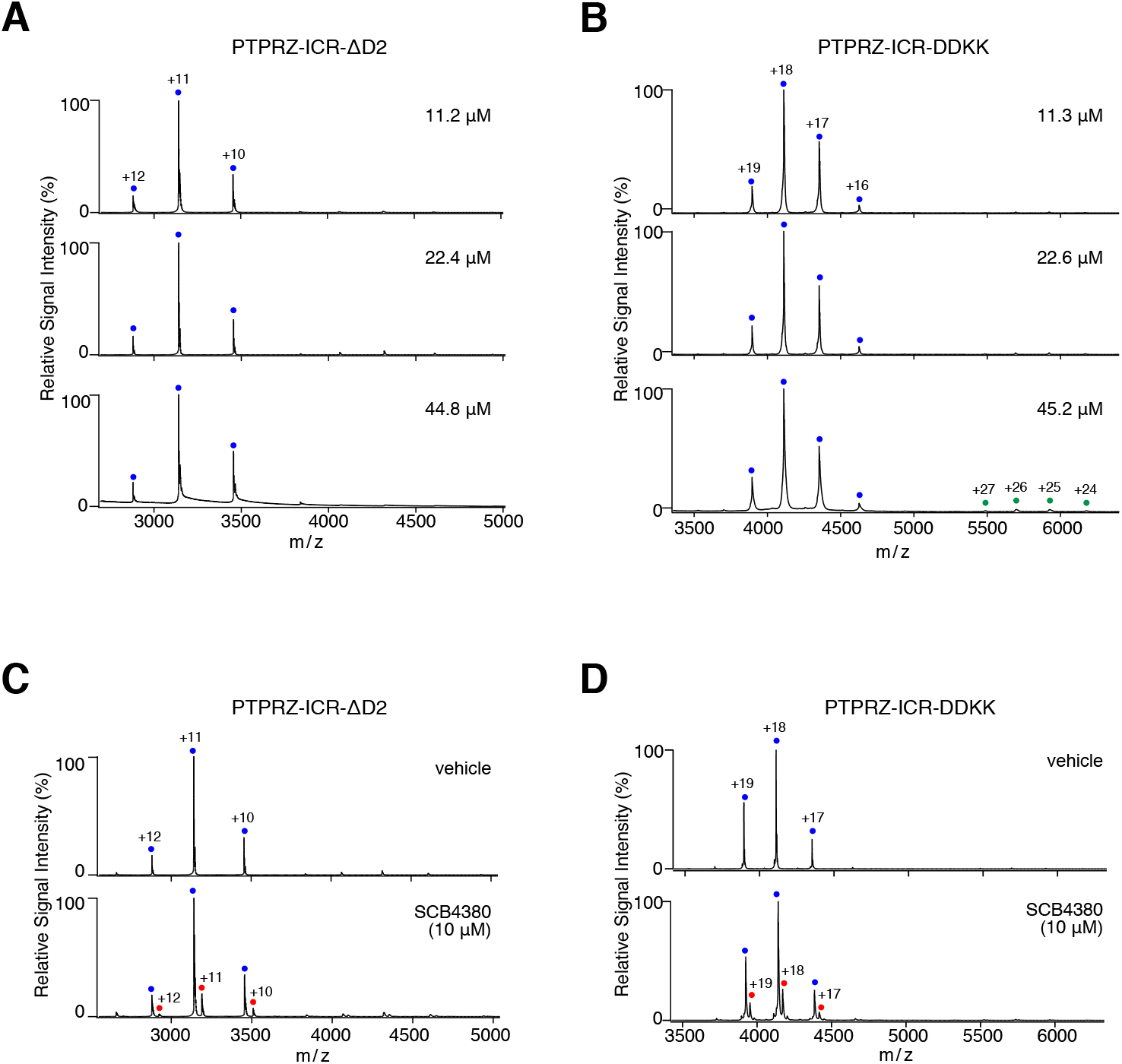
Monomer-dimer equilibrium of ΔD2 and DDKK mutants in aqueous solution. ***A,B***, The monomer-dimer equilibrium of ΔD2 (A) and DDKK (B) mutants of PTPRZ-ICR was analyzed as in Figure 3A. Product ion spectra derived from the corresponding monomers (ΔD2, 34,543 ± 0 Da; DDKK, 73,979 ± 0 Da) are indicated by blue dots with charge states. Peaks corresponding to ΔD2 or DDKK dimers were not observed. ***C,D***, ΔD2 (22.4 μM, *C*) and DDKK (22.6 μM, *D*) mutant proteins were mixed with SCB4380 at the indicated concentrations, and analyzed as in Figure 3B. Peaks corresponding to ΔD2 monomers complexed with SCB4380 (35,079 ± 9 Da) or DDKK monomers complexed with SCB4380 (74,523 ± 8 Da) are indicated by red dots with charge states. Masses were estimated from their parent ion peaks, and shown as the mean ± SD of three measurements.

**Figure 5.**
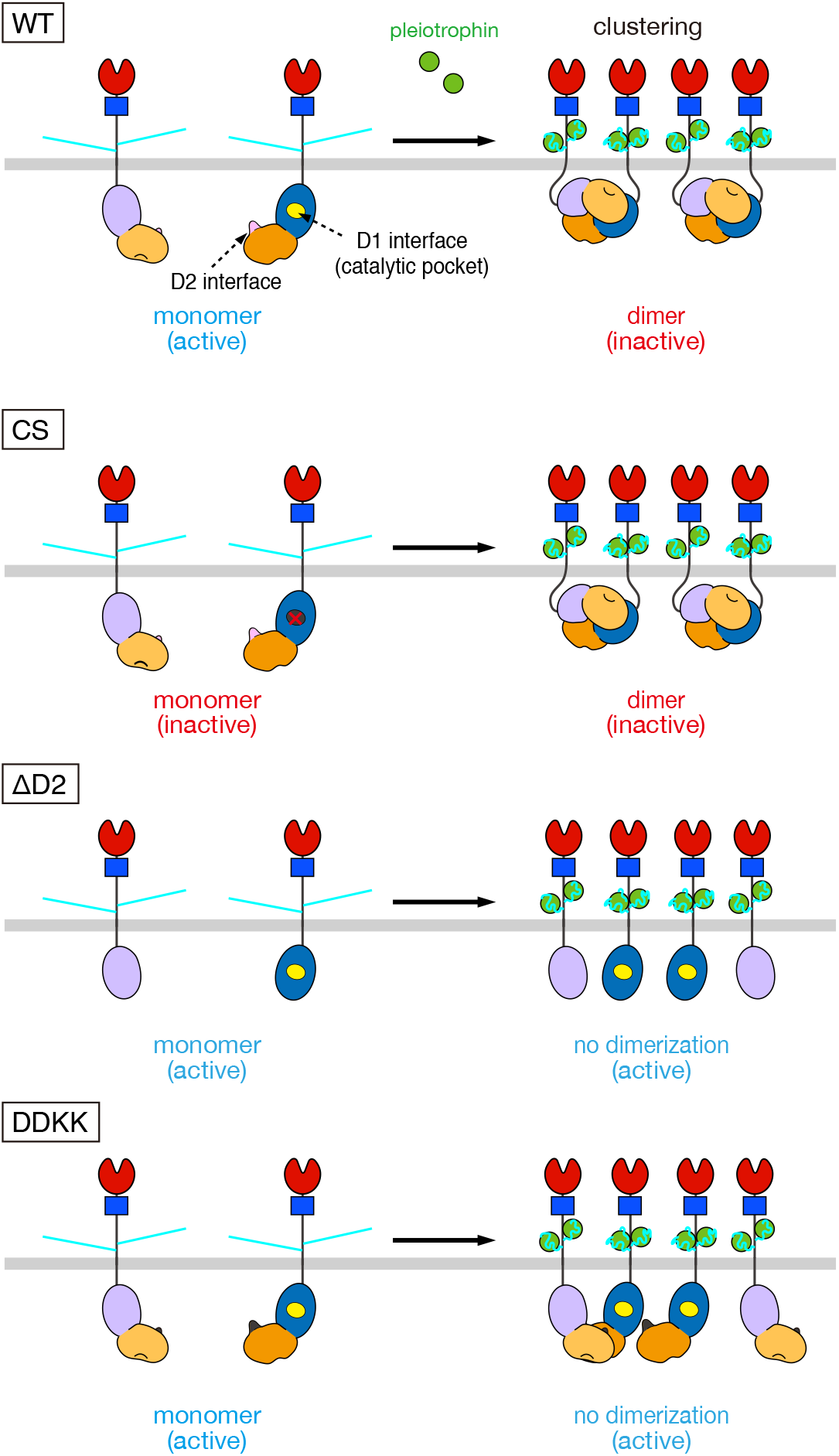
Models for ligand-induced PTPRZ clustering and dimerization. An experimental model of ligand-induced clustering for wild-type and mutant PTPRZ-B. PTPRZ-B receptors are expected to exist as monomers, but undergo clustering by extracellular ligand (PTN) binding, facilitating the “head-to-toe dimer” formation of the intracellular part, thereby masking the catalytic site of D1 by D2 from another (WT). On the other hand, the Cys-1930 to Ser mutation in the D1 domain (CS) abolished both catalytic activity and ligand-induced inactivation. The loss of the regulatory D2 domain (ΔD2) and the mutation in the dimer interface on D2 (DDKK) are catalytically active, but lacking the ligand-induced PTPase inactivation; however, ΔD2 and DDKK mutants are sensitive to PTPase inhibitors that bind to the active site. Domains are highlighted in different colors: carbonic anhydrase-like (CAH, red), fibronectin type III (FNIII, blue), PTP-D1 (blue or light purple), and PTP-D2 (orange or yellow-orange) domains. The extracellular regions of the three isoforms are highly glycosylated with chondroitin sulfate chains (light blue lines).

### Role of D2 in ligand-induced PTPRZ inactivation

We transfected four constructs expressing the wild-type PTPRZ-B receptor isoform (WT), PTPase-inactive mutant PTPRZ-B (CS), D2-deleted PTPRZ-B (ΔD2), or Asp-2178/2179 to Lys mutant (DDKK) into BHK-21 cells, which do not endogenously express PTPRZ proteins (17), together with p190RhoGAP (p190), one of the PTRPZ substrates. Similar to wild-type PTPRZ-B, ΔD2 and DDKK mutants reduced the tyrosine phosphorylation levels of FLAG-tagged p190, a key signaling molecule in the OPC differentiation process (13), significantly more than the mock control or PTPase-inactive CS mutant (Fig. S2). These results indicated that ΔD2 and DDKK mutants were active PTPase in living cells. The tyrosine phosphorylation level of p190 expectedly increased in wild-type PTPRZ-B-expressing cells upon the PTN treatment, but not in the other transfectants expressing the CS, DDKK, or ΔD2 mutant, or mock control (Fig. 6A). When these cells were treated with the cell-permeable PTPRZ inhibitor NAZ2329, p190 phosphorylation levels increased in cells with PTPRZ-B, or its mutant ΔD2 or DDKK, but not in cells with the CS mutant or mock control (Fig. 6B). These results supported the view that ΔD2 and DDKK mutations in PTPRZ-B receptors eliminated PTN sensitivity to inactivate PTPase because of a deficiency in dimerization.

**Figure 6.**
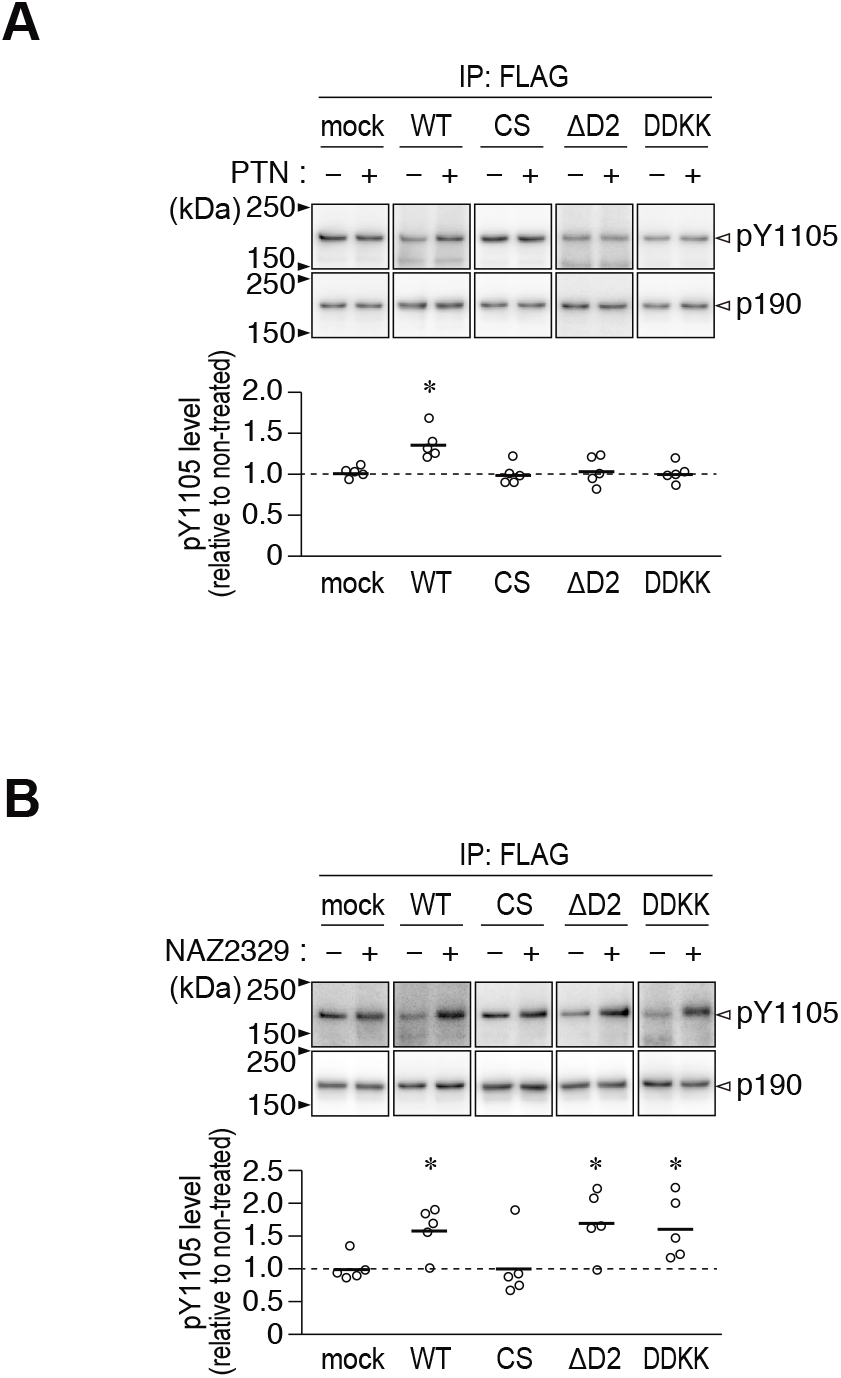
Effects of PTN and NAZ2329 on the tyrosine phosphorylation of p190RhoGAP in BHK-21 cells expressing PTPRZ or its mutant. ***A,B***, BHK-21 cells were transfected with wildtype PTPRZ-B, its CS, ΔD2, or DDKK mutant, or the mock control, together with FLAG-tagged p190RhoGAP. Transfected cells were treated with 100 nM PTN or vehicle (*A*), or 25 μM NAZ2329 or vehicle (*B*) for 1 h. The Tyr-1105 phosphorylation of FLAG-tagged p190RhoGAP immunoprecipitation was assessed using anti-Tyr(P)-1105 and anti-FLAG antibodies, respectively. The scatter plot shows Tyr-1105 phosphorylation levels relative to vehicle-treated levels (*n* = 5 independent cell culture per group). *, *P* < 0.05; significant different from the vehicle-treated control of each group by Welch’s t-test. Full-length blots for those that are cropped are shown in Supporting Information, Figure S4.

After the PTN treatment, PTPRZ-B receptors exhibit a patchy distribution in contrast to their diffuse distribution on the cell surface (11,17). The three mutants showed patchy distributions upon the PTN treatment as well as wild-type PTPRZ-B (compare “*con*” to “*PTN*” in Fig. 7), notwithstanding that their ICR proteins did not form a stable dimer *in vitro*. NAZ2329 did not affect the cell surface distribution of PTPRZ-B or its mutant forms (Fig. 7). These results indicated that the PTPRZ receptor clustering itself is independent of its PTPase inactivation.

**Figure 7.**
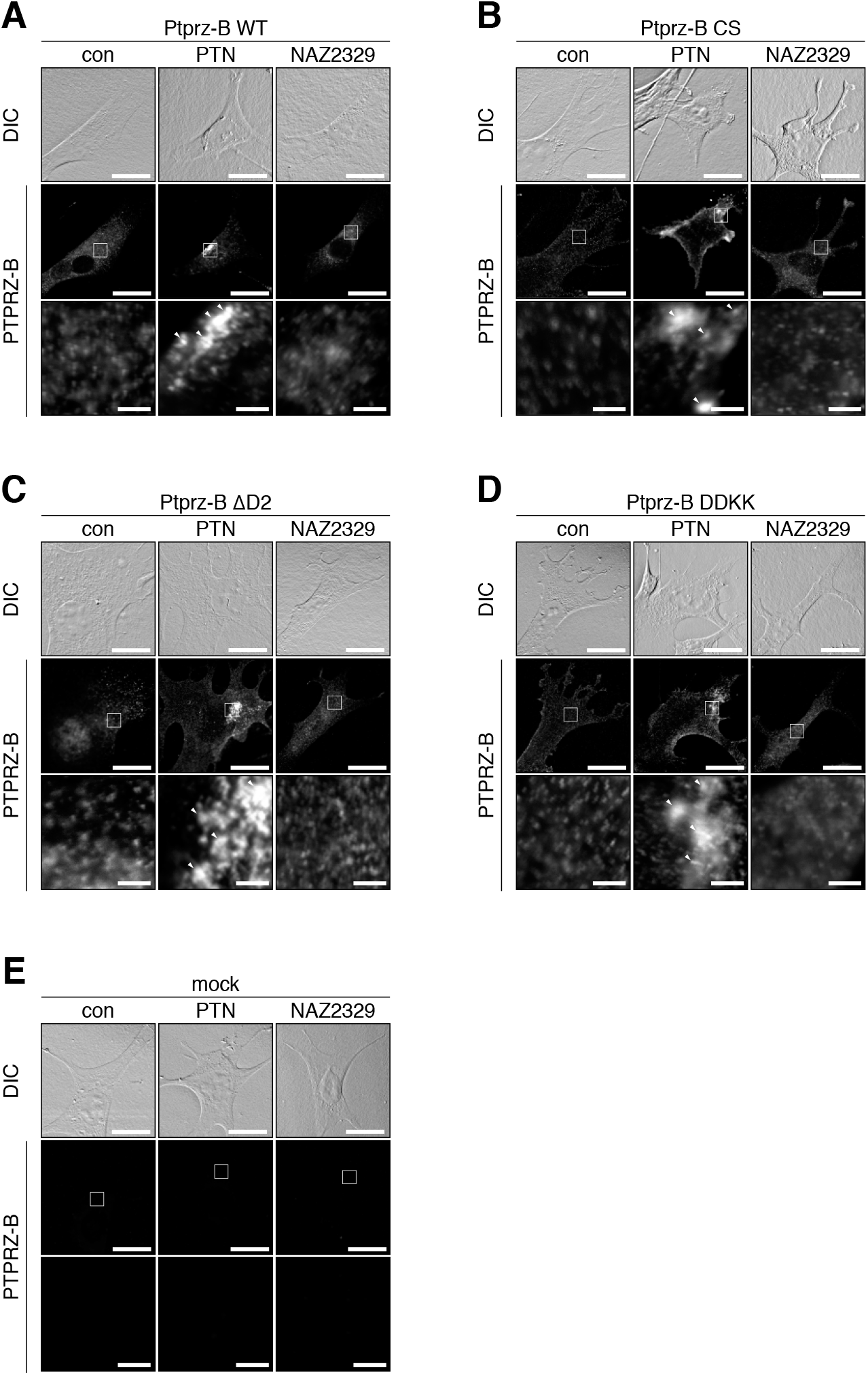
PTN-induced clustering of PTPRZ or its mutant in BHK-21 cells. ***A-E***, BHK-21 cells were transfected with wild-type PTPRZ-B (*A*), CS (*B*), ΔD2 (*C*), or DDKK (*D*) mutant, or the mock control (*E*), and treated with PTN or NAZ2329, as in Figure 6. Cells were fixed with formalin and stained with anti-PTPRZ-S against the extracellular epitope of PTPRZ. The membrane permeabilization step was omitted to prevent antibody staining of the cytosol, as described previously (17). The bottom picture in each column is an enlarged view of the rectangular region in the middle. Arrowheads, anti-PTPRZ-S-positive puncta. Scale bars, 100 μm (top and middle pictures) and 10 μm (bottom pictures), respectively. DIC, differential interference contrast.

### Effects of the ΔD2 mutation on PTN-enhanced OPC differentiation

To test its physiological significance, we performed a primary culture of neonatal mixed glial cells prepared from wild-type mice, *Ptprz*-knockout (KO) mice lacking all three PTPRZ isoforms (18), *Ptprz*-CS mutant knock-in mice (19), and ΔD2 mutant knock-in mice (Tanga N. et al., to be published elsewhere^5^). The cell number ratios of MBP-positive mature oligodendrocytes relative to NG2-positive OPCs were similar between wild-type and ΔD2 knock-in glial cells under normal differentiation conditions (without PTN, “*con*” in Fig. 8A,B). However, CS knock-in cells, as well as *Ptprz*-KO cells (13), showed higher differentiation levels than wild-type cells (Fig. 8A,B), indicating that PTPase activity negatively regulated OPC differentiation. We already reported that PTN enhanced OPC differentiation to MBP-positive cells in wildtype cells, but not in *Ptprz*-KO cells (13), and herein demonstrated that neither CS nor ΔD knock-in cells responded to the PTN stimulation (Fig. 8C). On the other hand, NAZ2329 enhanced the differentiation of ΔD2 knock-in cells as well as wild-type cells (20), but not *Ptprz*-KO cells or CS knock-in cells. These results indicated that “head-to-toe” RPTP dimerization is relevant to OPC differentiation as the molecular basis for PTN-induced PTPRZ inactivation.

**Figure 8.**
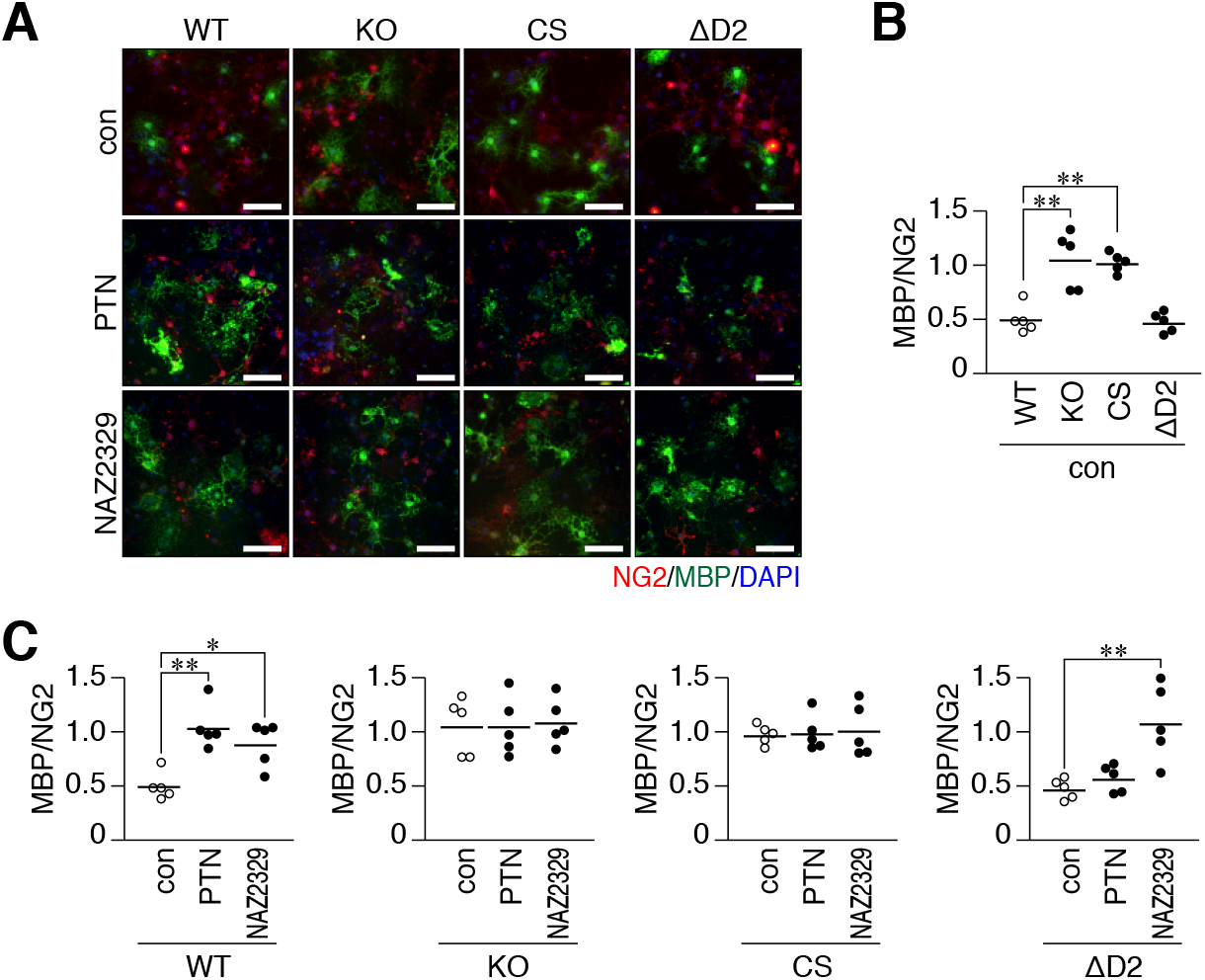
Effects of the *Ptprz*-CS or ΔD2 knock-in mutation on the PTN- or NAZ2329-stimulated differentiation of oligodendrocyte precursors cells into mature oligodendrocytes. ***A-C***, Glial cells were prepared from WT, KO, CS, or ΔD2 brains and cultured in differentiation medium in the presence of 100 nM PTN, 25 μM NAZ2329, or vehicle for 6 days. Fixed and permeabilized cells were stained with anti-NG2 proteoglycan (a marker of oligodendrocyte precursor cells, OPCs; *red*) and anti-MBP (oligodendrocytes, *green*) antibodies in conjunction with the DAPI labeling of nuclei (*blue*) (*A*). Scale bars, 100 μm. Scatter plots show the ratio of MBP-positive cells to NG2-positive cells, in which each circle corresponds to an independent cell culture (*n* = 5 each). *, *P* < 0.05; **, *P* < 0.01, significantly different from WT cells under control conditions (*B*), or the vehicle control of each group (*C*) by ANONA with Bonferroni’s *post hoc* tests.

## DISCUSSION

PTP family members contain a common CX5R(S/T) sequence, in that the cysteine residue that performs the initial nucleophilic attack of phospho-Tyr in substrate proteins is sensitive to oxidation and inactivation by reactive oxygen species (21). This redox-dependent inactivation is reversible (22,23), and is now widely recognized as a mechanism responsible for the down-regulation of PTPase activities in controlling signal outputs following an extracellular stimulus (21). On the other hand, the dimerization-induced inactivation of RPTP remains a controversial mechanism. The initial wedge model proposed from the crystal structure of PTPRZ-D1 is incompatible with the tandem PTP domain structures due to a steric clash with the D2 domain (7,8). The “head-to-toe dimer” inactivation model was subsequently proposed from the crystal structure of PTPRG-D1D2 segment, with D2 masking the catalytic site of D1 (8); however, its validity as the inhibitory regulation mechanism has not yet been investigated. The crystal structure of the D1D2 segment of PTPRZ also showed the “head-to-toe dimer” conformation (Figs. 1 and 2). Consistent with this structure, the inhibitory effects of PTN were abolished in living cells by the deletion or mutation of the regulatory D2 domain (Figs. 6 and 8), confirming its physiological relevance. To the best of our knowledge, the present study is the first to elucidate the molecular basis for the PTN-induced inactivation of PTPase.

Previous findings obtained from analytical ultracentrifugation for PTPRG indicated the concentration-dependent and stable dimerization of the D1D2 segment in solution with a K*d* value of ~4 μM (8). Our results for PTPRZ showed that the peak heights of monomer and dimer peaks were similar when applied at 25 μM in native MS analyses (Fig. 3A), estimating an apparent K*d* of ~7 μM that appears to be close to that of PTPRG. On the other hand, the competitive inhibitor, SCB4380 dose-dependently reduced dimer peaks (Fig. 3B). These results suggest that PTPRZ-ICR proteins exist in dynamic monomer-dimer equilibrium in solution. It is important to note the close proximity of Asp-1845 on one PTPRZ molecule and Glu-1848 on the other (Fig. 2Ac), which may cause charge repulsion and prevent stable dimer formation. The rapid association/dissociation of intracellular D1D2 may be a suitable property for a regulatory mechanism.

D2 truncation (ΔD2) abolished the dimerization abilities of PTPRZ-ICR proteins without changing SCB4380 binding abilities *in vitro* (Fig. 4). In living cells, the ΔD2 mutant receptors of PTPRZ-B exogenously expressed were not inactivated by PTN (Fig. 6), indicating a causal relationship between the masking effects of D2 on the catalytic site and ligand-induced PTPase inactivation. The results of PDBePISA showed multiple interactions in the dimer interface between D1 and D2 (Table S2). The interactions between acidic Asp-2178 and Asp-2179 residues in D2 and basic Lys-1837 and Arg-1940 residues in D1 are expected to be major stabilizing forces of the dimer structure. The substitution of Asp-2178 and Asp-2179 with Lys residues (DDKK mutant) largely abolished dimerization abilities similar to the D2 deletion (Fig. 4). The DDKK substitution also disrupted the dimerization ability of PTPRG (8). Therefore, “head-to-toe dimerization” may be a conserved regulatory mechanism in two members of the R5 RPTP subfamily.

In contrast to the close similarities between the intracellular parts in PTPRZ and PTPRG, their extracellular regions display different characteristic features. The most obvious difference is the specific modification of PTPRZ with extensive glycosaminoglycans (9,24), which is of crucial importance for the ligand-dependent regulation of PTPase inactivation (11). The negatively-charged chondroitin sulfate moiety is essential for the diffuse distribution of PTPRZ receptors at the cell membrane due to electrostatic repulsion, thereby maintaining them in a catalytically active monomer state (11). Notably, chondroitin sulfate chains are required for high-affinity ligand binding (25,26). The accumulative binding of positively charged PTN attenuates electrostatic repulsion between chondroitin sulfate chains and induces the clustering of PTPRZ receptors (11), which may be a prerequisite for allowing the “head-to-toe dimerization” of intracellular parts in living cells (Figs. 6 and 8). On the other hand, these chondroitin sulfate modifications were not found in PTPRG (27), and no ligand molecules have been identified. Thus, regarding PTPRG, studies to confirm the *in vivo* relevance of this inhibitory regulation mechanism will be necessary in the future.

Barr AJ et al. also reported that the orientations of D1 and D2 of PTPRC, PTPRE, PTPRF, PTPRS, and PTPRG were highly conserved (8), suggesting that “head-to-toe dimerization” is a general regulatory mechanism for these RPTPs. Most of the residues involved in the dimer interface of the D1 side are well conserved not only in tandem-type RPTP members, but also in all PTP family members, whereas key Asp-Asp residues on the D2 side of R5 members (PTPRZ and PTPRG) are changed to Arg-Lys in the R1/R6 member (PTPRC), Gln-Asp/Glu in R2B (PTPRK, PTPRM, PTPRT, and PTPRU), Asp-Gly in R2A members (PTPRD, PTPRF, and PTPRS), or Glu-Asn in a R4 member (PTPRA) (see Fig. S3). These changes may abolish or reduce dimer formation. As one exception, only PTPRE in the R4 subfamily contains acidic Glu-Glu residues as R5 RPTP members. However, dimer formation was not found in PTPRE-D1D2, similar to PTPRA, PTPRC, and PTPRM (8). Regarding R4 members, acidic Glu residue(s) are adjacent to conserved Lys (corresponding to Lys-1837 of rat PTPRZ) on the D1 side, which may interfere with dimerization.

Besides homodimerization, the heterodimerization of RPTP family members may be possible *in vivo*. In support of this view, the D2 domains of PTPRZ, PTPRA and PTPRM were isolated in a yeast two-hybrid screen as interactors of the cytoplasmic region of PTPRZ (28). The dimerization of RPTPs may also be regulated by post-translational modifications, such as oxidation (21,29,30). The involvement of these factors in “head-to-toe dimerization” needs to be investigated in future studies.

## EXPERIMENTAL PROCEDURES

### Ethics statement and experimental animals

All animal experimental protocols used in the present study were approved by the Institutional Animal Care and Use Committee of National Institutes of Natural Sciences, Japan (Approval number, 17A020). Mouse pups were handled gently to minimize stress and quickly decapitated without anesthesia, and brains were collected.

*Ptprz*-null knockout mice (18) and Ptprz-ΔD2-knockin mice^5^ and *Ptprz*-CS-knockin mice (19) were backcrossed with the inbred C57BL/6J strain (CLEA Japan) (wildtype) for more than ten generations.

### Reagents and antibodies

SCB4380 (31), NAZ2329 (20), and recombinant PTN (32) were described previously. Stock solutions of PTN were prepared as 100 μg/ml with PBS (4.3 mM Na_2_HPO_4_, 1.4 mM KH_2_PO_4_, 137 mM NaCl, and 2.7 mM KCl at pH 7.4) containing 100 μg/ml BSA and stored at −85°C until used. Anti-PTPRZ-S, rabbit polyclonal antibodies against the extracellular region of PTPRZ (33), and anti-pY1105, purified rabbit polyclonal antibodies against phosphorylated Tyr-1105 of p190RhoGAP (34) were described previously. The following are the specificities and sources of the commercially available antibodies used in the present study: antimyelin basic protein (catalog #sc-13914, Santa Cruz Biotechnology), anti-NG2 proteoglycan (catalog #AB5320, Millipore), anti-p190RhoGAP (catalog #610150, BD Biosciences; and catalog #12164, Cell Signaling Technology), anti-phosphotyrosine (PY20; catalog #ab16389, Abcam), and anti-FLAG (catalog #F7425 for Western blotting and #F3165 for immunoprecipitation; Sigma-Aldrich).

### X-ray crystallography of rat PTPRZ-D1D2

Recombinant proteins of rat PTPRZ-D1D2 (residues 1699-2316) were expressed using a baculovirus-silkworm expression system, and purified as described (31). The initial screening of crystallization conditions was performed using PACT (Molecular Dimensions). Crystals were obtained at 277K in 14% (*w/v*) PEG3350, 200 mM potassium fluoride, and 0.1 M bis-Tris propane, pH 6.5 by the sitting-drop vapor diffusion method. After brief soaking in 14% PEG3350, 3.6 M potassium fluoride, and 0.1 M bis-Tris propane, pH 6.5, crystals were flash-frozen in a cold nitrogen stream at 100 K. X-ray diffraction data were collected using an RAXIS IV^++^ imaging-plate area detector mounted on a Rigaku MicroMax-007 rotating-anode source with CuKα radiation (*λ* = 1.5418 Å, 40 kV, 20 mA). Diffraction data were processed and scaled with HKL2000 (35). The crystal structure was solved by the molecular replacement method using the structure of the D1-D2 domain of PTPRG (PDB ID, 2NLK). A molecular-replacement calculation was performed using AMoRe (36). The structural model was manually fit using Coot (37) and refined with Refmac-5 (38). Interface areas and interactions were calculated by “Protein interfaces, surfaces and assemblies” service PISA (39) at the European Bioinformatics Institute. Figures were created by Pymol (DeLano Scientific).

### Mass spectrometry under nondenaturing conditions

Purified PTPRZ-D1D2 (31), PTPRZ-ΔD2 (residues 1699-1997), and PTPRZ-D1D2-DDKK mutant proteins expressed using the baculovirus-silkworm expression system were buffer exchanged to 100 mM ammonium acetate, pH 7.0 by passing through a MicroBioSpin-6 column (Bio-Rad), diluted 100-fold with the same buffer, and then kept on ice. Ten-microliter aliquots were mixed with equal volumes of individual fining compound solutions, and samples (~2 μl for each analysis) were immediately analyzed by nanoflow electrospray using in-house gold-coated glass capillaries. Spectra were recorded on a SYNAPT G2-Si HDMS mass spectrometer (Waters) in the positive ionization mode at 1.36 kV with 150 V of a sampling cone voltage and source offset voltage, 0 V of trap and transfer collision energy, and 2 ml/min trap gas flow. Spectra were calibrated using 1 mg/ml cesium iodide and analyzed by MassLynx software (Waters).

### Expression plasmids

The mammalian expression plasmids, pZeoPTPζ (for PTPRZ-B) (40) and pFLAG-p190 (for FLAG-tagged p190RhoGAP), were described previously (41), and pZeoPTPζ-ΔD2 (the D2 deletion mutant of PTPRZ-B) was generated using pZeoPTPζ as a template with a Quikchange multisite-directed mutagenesis kit (Stratagene).

### BHK-21 cell culture and DNA transfection

Hamster kidney BHK-21 cells have been maintained in our laboratory. BHK-21 cells were cultured in DMEM (Life Technologies) supplemented with 10% fetal bovine serum (FBS, Nichirei Biosciences) in a humidified incubator at 37°C with 5% CO_2_. DNA transfection was performed using Lipofectamine 2000 reagent (Thermo Fisher Scientific) according to the manufacturer’s directions.

### Primary mixed glial culture

A primary mixed glial cell culture was performed as described previously (13). Cortex tissues obtained from mouse brains on postnatal day 1 were dissociated with papain (Worthington Biochemical). Dissociated cells (2.0 × 10^4^ cells) were cultured on a poly-L-ornithine-coated 35-mm dish with DMEM differentiation medium mixed 1:1 (*v*/*v*) with Ham’s F-12 (DMEM/F-12, Life Technologies), 1× GlutaMAX (Life Technologies), 1× N2 supplement (Life Technologies), 10 μg/ml of the AA homodimeric form of platelet-derived growth factor (PDGF-AA, Wako Pure Chemical), 0.5% FBS, 100 μg/ml of bovine serum albumin (BSA, Sigma-Aldrich), 10 nM biotin (Sigma-Aldrich), and 30 ng/ml thyronine/thyroxine (Sigma-Aldrich) in a humidified incubator at 37°C with 5% CO_2_. On the sixth day of culture, cells were fixed and stained with anti-NG2 (a specific marker for OPCs) and anti-MBP (a marker for matured oligodendrocytes). Differentiation from OPC to oligodendrocytes was estimated from the ratio of MBP-positive cells to NG2-positive cells.

### Immunocytofluorescence staining

Cells were fixed with 4% paraformaldehyde in PBS for 30 min. Fixed cells were permeabilized and blocked with 4% non-fat dry milk and 0.1% Triton X-100 in TBS (10 mM Tris-HCl, pH7.4, 150 mM NaCl) for 30 min, and then incubated overnight with the respective primary antibodies at 4°C. When cell surface proteins were analyzed, the membrane permeabilization step was omitted to prevent antibody staining of the cytosol, as described previously (17). Bound primary antibodies were visualized with Alexa Fluor-conjugated secondary antibodies (Life Technologies) or DyLight Amine-Reactive Dyes (DyLight 488 NHS Ester, Thermo Fisher Scientific) according to the standard procedure. Digital photomicrographs of individual specimens were taken with the Biozero BZ-8000 (Keyence), or LSM 700 confocal microscope (Zeiss).

### Protein extraction and chABC digestion

Proteins were extracted from mouse brains or cultured cells with 1% Nonidet P-40 in TBS containing 1 mM vanadate, 10 mM NaF, and protease inhibitors (EDTA-free complete, Roche Molecular Biochemicals). Regarding the detection of PTPRZ proteins, tissue extracts with 1% Nonidet P-40 in TBS containing protease inhibitors (Complete, Roche Molecular Biochemicals) were mixed with an equal volume of 0.2 M Tris-HCl, 60 mM sodium acetate, 10 mM EDTA, pH 7.5 containing 25 micro-units/μl of chABC (Sigma), or not (control), at 37°C for 1 h.

### Immunoprecipitation and Western blotting

Protein extracts were precleaned with Protein G Sepharose (GE Healthcare) and subjected to immunoprecipitation with a combination of anti-FLAG M2 and Protein G Sepharose beads. Immunocomplexes were then separated by SDS-PAGE, followed by semi-dry electroblotting onto a polyvinylidene difluoride membrane. After blocking with 4% non-fat dry milk and 0.1% Triton X-100 in TBS, membranes were incubated overnight with the respective antibodies. The binding of antibodies was detected with Luminata Forte Western HRP Substrate (Millipore). Regarding the detection of tyrosine phosphorylated proteins in Western blotting, antibodies (PY20 or anti-pY1105) were diluted with blocking solution (1% BSA and 0.1% Triton X-100 in TBS).

### Image and statistical analyses

Quantitative image analyses were performed using ImageJ software (NIH) or Adobe Photoshop CS6 software (Adobe Systems). Statistical analyses were performed using IBM SPSS Statistics 25 software (IBM).

## Acknowledgments

We thank Ms. Yoshiko Isoshima and Ms. Norie Nakanishi for their technical assistance, and Ms. Akiko Kodama for her secretarial assistance. Immunofluorescence photomicrograph images were acquired at the Spectrography and Bioimaging Facility, NIBB Core Research Facilities. Native MS experiments were supported by Joint Research by Exploratory Research Center on Life and Living Systems (ExCELLS).

## Author contributions

A.F. and M.N. designed the research. H.S. performed X-ray crystal structure analyses. K.I., A.F., and S.U. performed mass analyses under non-denaturing conditions. N.T., and R.S. generated mutant plasmids. A.F. performed enzyme assays. N.T., K.K., R.S., and A.F. performed cell culture experiments. K.K. performed primary culture experiments. N.T., K.K., and A.F. performed statistical analyses and prepared figures. A.F. and M.N. wrote the manuscript.

## FOOTNOTES

This work was supported in part by the Japan Society for the Promotion of Science (JSPS) KAKENHI Grant Numbers 17K07069 and 26110722 (to A.F.), and 17K07355 (to K.K.).

## Abbreviations

PTPRZ: protein tyrosine phosphatase receptor type Z
CS: chondroitin sulfate
PTN: pleiotrophin
OPC: oligodendrocyte precursor cell
CNS: central nervous system
GAP: GTPase-activating protein
RPTP: receptor-type protein tyrosine phosphatase
MBP: myelin basic protein
ECM: extracellular matrix
HB-GAM: heparin-binding growth-associated molecule
MK: midkine
IL-34: interleukin-34
NG2: neural/glial antigen 2 chondroitin sulfate
chABC: chondroitinase ABC
GFAP: glial fibrillary acidic protein
GlcA: glucuronic acid
GalNAc: N-acetylgalactosamine
MS: multiple sclerosis.

## SUPPORTING INFORMATION

**Table S1.** Data collection and refinement statistics of the X-ray structure of rat PTPRZ-D1D2.

**Figure S4. Full-length blots for Figure 6.**
| Data collection and refinement statistics for<br>the PTPRZ-ICR (D1+D2) |  |  |
| --- | --- | --- |
| **Data Collection** |  |  |
| Space group |  | *P*2<sub>1</sub>2<sub>1</sub>2 |
| Cell Dimensions |  | 53.78 |
| *a,b,c* (Å) |  | 127.73, 147.59, 65.68 |
| Resolution (Å) |  | 3.32 |
| Redundancy |  | 3.3 (2.7) |
| Completeness (%) |  | 84.0 (84.0) |
| I/sigma |  | 13.0 (3.3) |
| *R*<sub>merge</sub> |  | 6.2 (22.2) |
| **Refinement** |  |  |
| Resolution (Å) |  | 3.32 |
| No. of reflections |  | 14903 |
| *R*<sub>work</sub> / *R*<sub>free</sub> |  | 23.9/31.8 |
| No. of atoms |  |  |
| Protein |  | 8548 |
| B-factors |  |  |
| Protein |  | 85.05 |
| R.m.s deviations |  |  |
| Bond lengths (Å) |  | 0.010 |
| Bond angles (°) |  | 1.491 |

**Table S2.** Summary of interaction residues.

| Hydrogen bonds |  |  |  | Salt bridges |  |  |  |
| --- | --- | --- | --- | --- | --- | --- | --- |
| no. | chain A | distance (Å) | chain B | no. | chain A | distance (Å) | chain B |
| 1 | R1835 [NH2] | 2.58 | W2112 [O ] | 1 | K1837 [NZ ] | 3.43 | D2178 [OD2] |
| 2 | K1837 [NZ ] | 3.43 | D2178 [OD2] | 2 | K1837 [NZ ] | 3.51 | D2179 [OD1] |
| 3 | K1837 [NZ ] | 3.51 | D2179 [OD1] | 3 | R1940 [NE ] | 3.99 | D2178 [OD1] |
| 4 | G1904 [N ] | 2.81 | Q2177 [O ] | 4 | R1940 [NH2] | 3.58 | D2178 [OD1] |
| 5 | A1936 [N ] | 2.94 | D2179 [OD2] | 5 | D2178 [OD2] | 3.55 | K1837 [NZ ] |
| 6 | R1940 [NH2] | 3.58 | D2178 [OD1] | 6 | D2179 [OD1] | 3.56 | K1837 [NZ ] |
| 7 | Q1978 [NE2] | 2.79 | D2178 [O ] | 7 | D2178 [OD1] | 3.84 | R1940 [NE ] |
| 8 | E2174 [OE1] | 2.93 | N1759 [ND2] | 8 | D2178 [OD1] | 3.65 | R1940 [NH2] |
| 9 | W2112 [O ] | 3.45 | R1835 [NH2] |  |  |  |  |
| 10 | D2178 [OD2] | 3.55 | K1837 [NZ ] |  |  |  |  |
| 11 | D2179 [OD1] | 3.56 | K1837 [NZ ] |  |  |  |  |
| 12 | Q2177 [O ] | 3.09 | G1904 [N ] |  |  |  |  |
| 13 | D2179 [OD2] | 2.89 | A1936 [N ] |  |  |  |  |
| 14 | D2178 [OD1] | 3.84 | R1940 [NE ] |  |  |  |  |
| 15 | D2178 [OD1] | 3.65 | R1940 [NH2] |  |  |  |  |
| 16 | D2178 [O ] | 3.25 | Q1978 [NE2] |  |  |  |  |

note: D1 (blue), D2 (orange)

**Figure S1.**
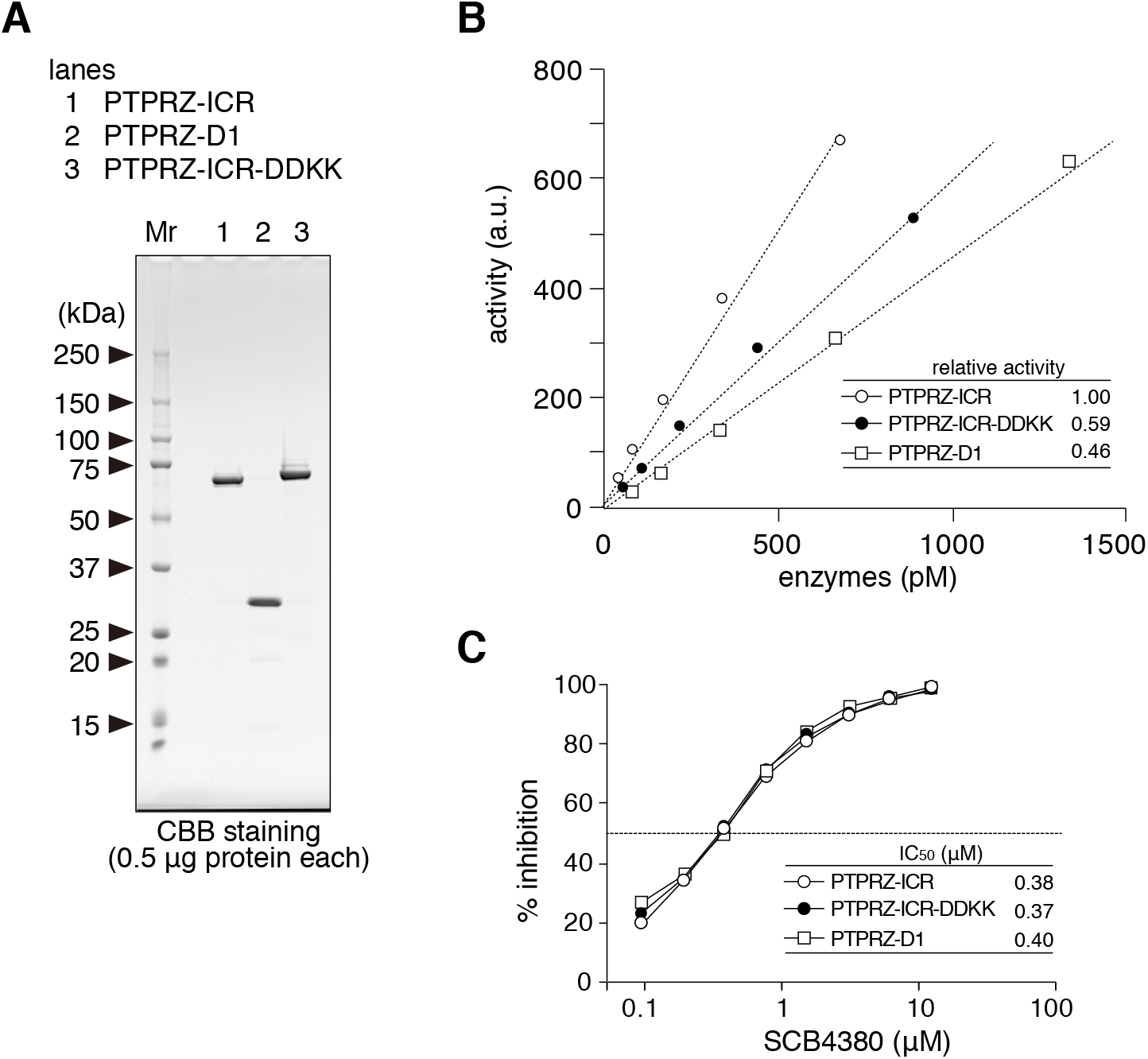
Catalytic activity and IC_50_ values of recombinant PTPRZ proteins. ***A***, The purities of recombinant PTPRZ proteins were analyzed by SDS-PAGE and CBB staining. ***B***, The catalytic activities of recombinant enzymes were measured using fluorogenic DiFMUP (non-specific PTP substrate, 6,8-difluoro-4-methylumbiliferyl phosphate) as described previously (31). ***C***, Concentration-inhibition curves of SCB4380 for the hydrolysis of DiFMUP were analyzed as described previously (31). The IC_50_ value of rat PTPRZ-D1D2 (0.38 μM) was similar to that of the human PTPRZ enzyme (0.40 μM) reported previously (31).

**Figure S2.**
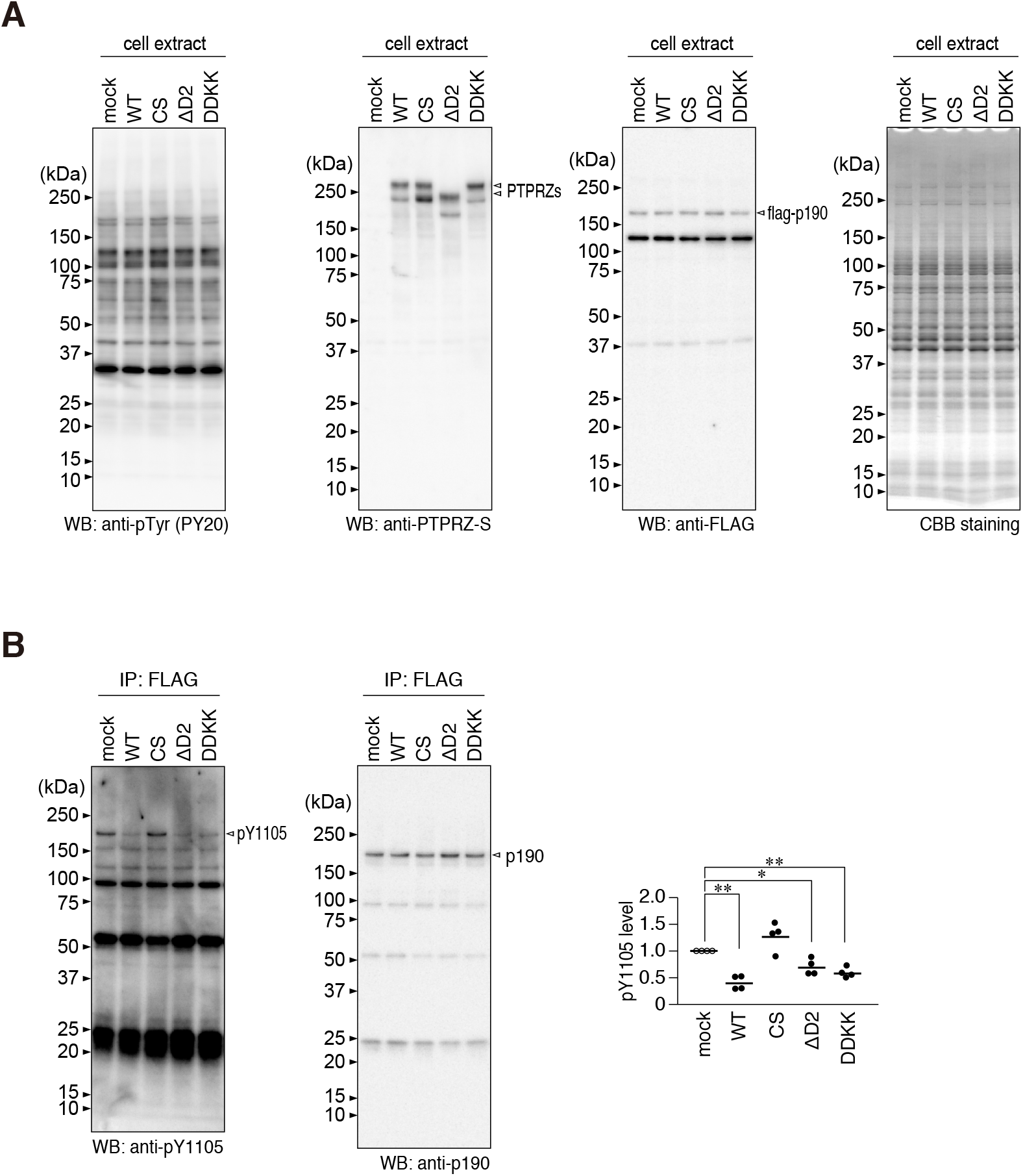
Expression of PTPRZ-B and its mutants in BHK-21 cells. ***A***, BHK-21 cells were transfected with wild-type PTPRZ-B, CS, ΔD2, or DDKK mutant, or the mock control, together with FLAG-tagged p190RhoGAP. Cellular proteins were extracted with 1% NP-40 buffer. The overall tyrosine phosphorylation pattern of cellular proteins and the expression of each protein were assessed by Western blotting using anti-phosphotyrosine (PY20 for the detection of the total tyrosine phosphorylation of cellular proteins), anti-PTPRZ-S (for PTPRZ-B and its mutant proteins), and anti-FLAG antibodies (for FLAG-tagged p190 RhoGAP). Protein amounts were verified by CBB staining. ***B***, The Tyr-1105 phosphorylation of FLAG-tagged p190 RhoGAP immunoprecipitation was assessed using anti-Tyr(P)-1105 and anti-FLAG antibodies. The scatter plot shows Tyr-1105 phosphorylation levels relative to those in the mock control (*n* = 4 independent cell culture per group). *, *P* < 0.05, significant different from the vehicle-treated control of each group by Welch’s *t*-test.

**Figure S3.**
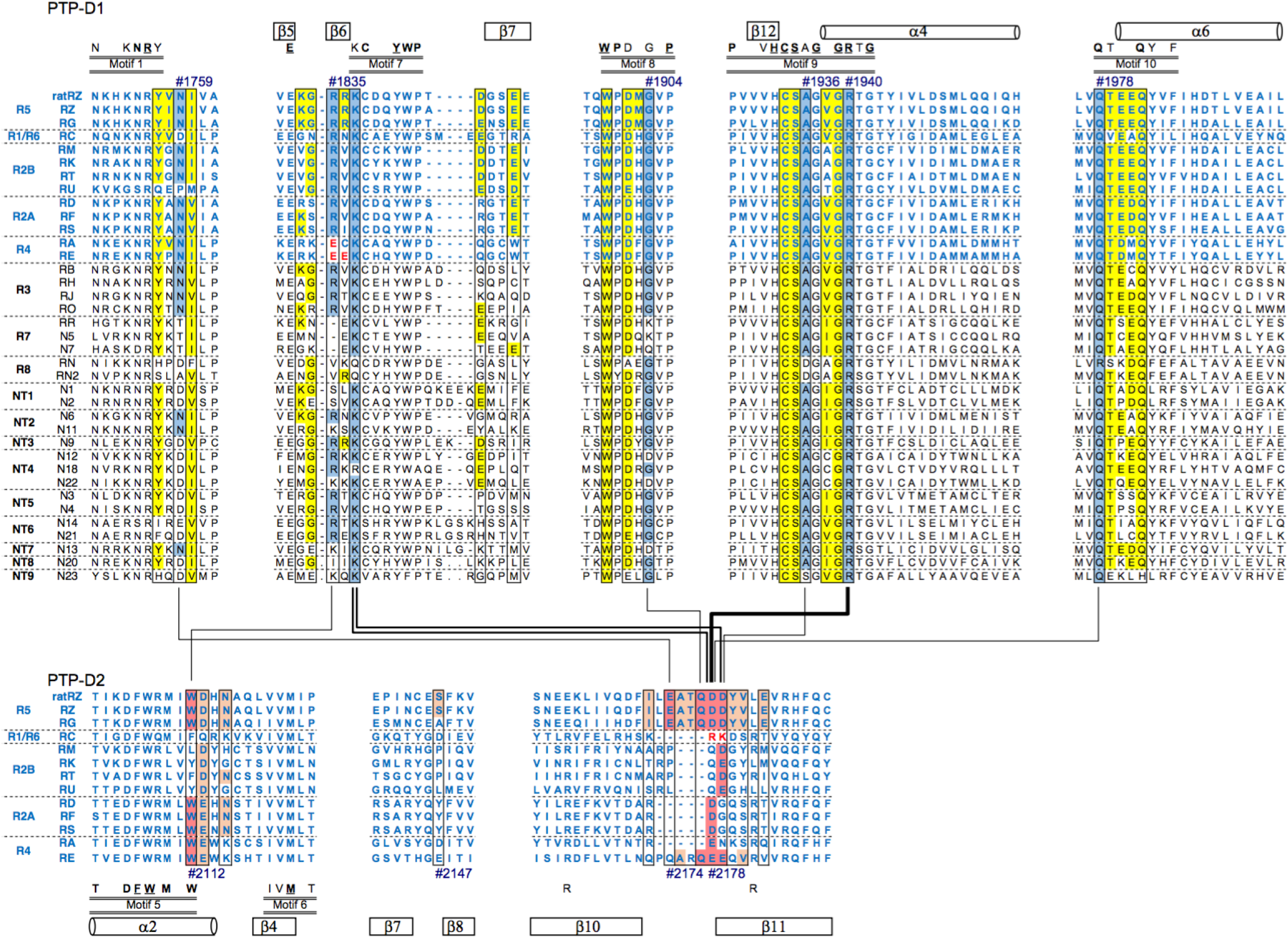
Sequence comparison of the dimer interface in the RPTP subfamily. Sequence alignment in the dimer interface of PTP-D1 and PTP-D2 domains of rat PTPRZ (rat RZ) and 37 human members of the classical PTP family: 8 receptor-type transmembrane subtypes (R1/6 to R7) and nine non-transmembrane subtypes (NT1 to NT9). In RPTPs with tandem domains, the membrane proximal D1 domain exhibits catalytic activity, while the distal D2 domain is either inactive or has negligible (RA and RE) catalytic activity (3). Upper and lower alignments show the sequence alignments of PTP-D1 and PTP-D2, respectively. Amino acid numbers correspond to the residues involved in dimerization in rat PTPRZ, and their interactions are represented by red lines (see Table S2). The residues forming the dimerization surface are boxed with black lines (see text and Fig. 2), and their residues are colored according to common characteristics: hydrophobic (black), aromatic (gray), basic (blue), acidic (red), and polar (N/Q, pink; S/T, magenta) residues. The locations of α-helices and β-strands and PTP signature motifs shown at the top were adopted from Andersen *et al*. (3).

**Figure S4.**
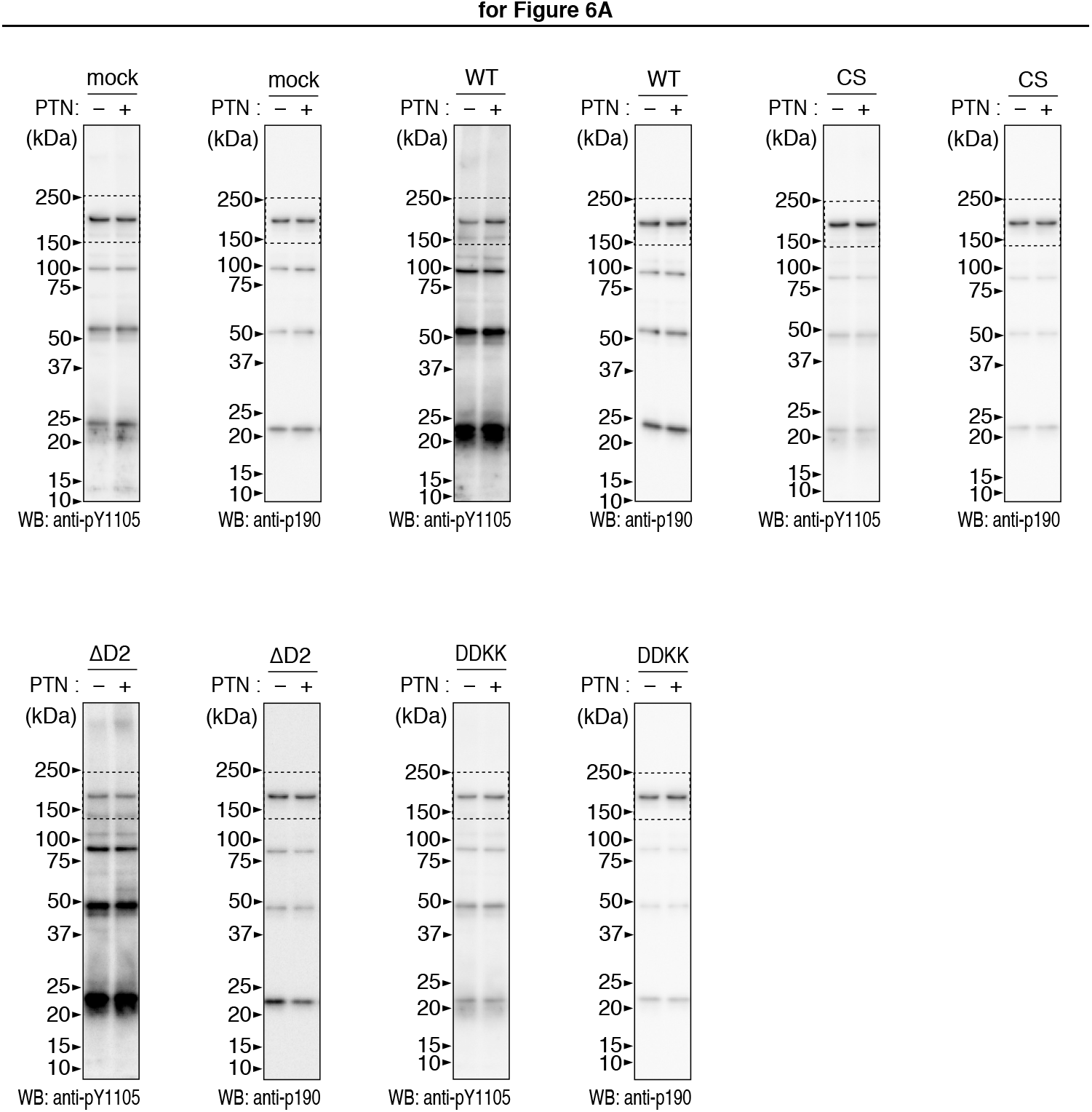
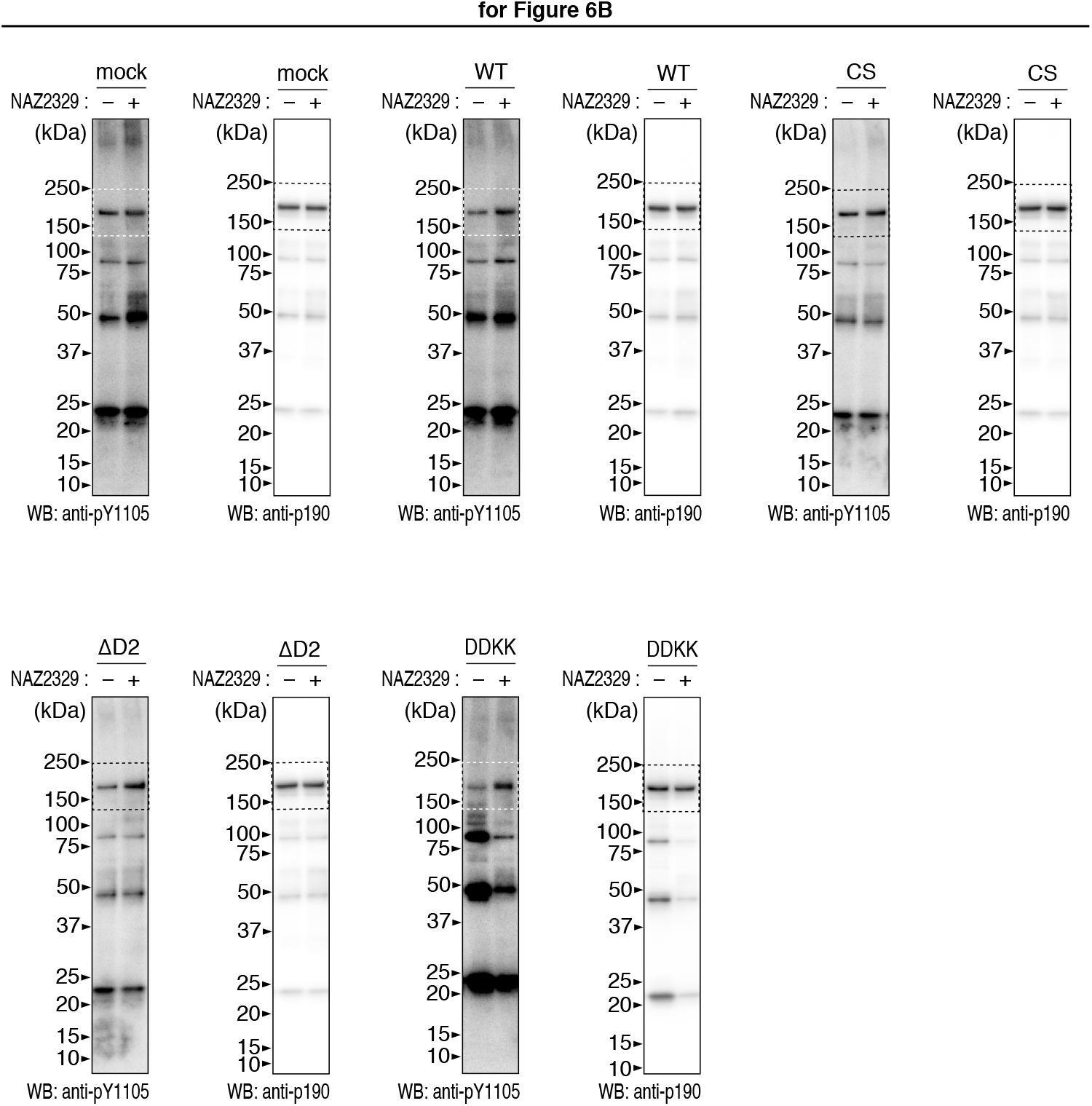
Full-length blots for Figure 6.

## Footnotes

4 A. Fujikawa, K. Kuboyama, and M. Noda, unpublished data.

5 N. Tanga, A. Fujikawa, M. Noda, et al., submitted for publication.

